# The accessory type III secretion system effectors shape intestinal inflammatory infection outcomes

**DOI:** 10.1101/2025.04.15.648969

**Authors:** Priyanka Biswas, Julia Sanchez-Garrido, Zuzanna Kozik, Vishwas Mishra, David Ruano-Gallego, Rita Berkachy, Sarah Jordan, Joshua L.C. Wong, Jyoti S. Choudhary, Gad Frankel

## Abstract

Injection of effectors via a type III secretion system (T3SS) is an infection strategy shared by various Gram-negative bacterial pathogens, many infecting mucosal surfaces. While individual T3SS effectors are well characterized, their network-level organization and the distinction between core and accessory effectors remain incompletely understood. Here, by systematically dissecting the T3SS effector network of the enteric mouse pathogen *Citrobacter rodentium* (CR) we identified a subset of 12 accessory effectors that, while dispensable for colonization, significantly alter infection outcomes. A strain lacking the accessory effectors (CR_M12_) remained virulent in susceptible mouse hosts yet resulted in reduced epithelial barrier damage, inflammation, and immune cell infiltration in resistant mice. Deep proteomic analysis specifically targeting CR-attached colonic epithelial cells revealed that, despite lacking 39% of its effector repertoire, infection with CR_M12_ results in similar changes to global protein expression as seen in mice infected with the wild-type strain, though key regulators of barrier integrity were differentially expressed. Using a host model with impaired barrier repair, we confirmed that accessory effectors shape infection outcomes without significantly impacting virulence. This study refines the concept of core and accessory effectors, providing a basis for further studies into effector-driven host adaptation.

## Introduction

Many Gram-negative bacterial pathogens infecting humans, animals, and plants execute their infection strategies via injection of type III secretion system (T3SS) effectors into the infected eukaryotic cells. The T3SS apparatus and its cognate effectors, which are frequently encoded in pathogenicity islands, prophages and/or plasmids, are found in pathogens like Enteropathogenic *Escherichia coli* (EPEC), Enterohemorrhagic *E. coli* (EHEC), *Citrobacter rodentium* (CR), *Salmonella enterica*, *Shigella spp.*, *Pseudomonas spp.*, and *Yersinia spp*^1,2,3^. While the T3SS injectisome is structurally conserved across these pathogens, the number and function of the effectors change dramatically from one pathogen to the other. Moreover, the effector composition varies considerably even among closely related pathotypes. For instance, *Salmonella enterica* serovar Typhimurium encodes twice as many effectors as *S. enterica* serovar Typhi^4,5^, while clinical EPEC isolates possess between 21 and 40 effectors^6^. Despite this variation, EPEC strains induce diarrheal diseases, characterized by effacement of the brush border microvilli, intimate bacterial attachment to the apical surface of intestinal epithelial cells (IECs), and the formation of actin-rich pedestals at the host-pathogen interface^7,8^.

CR is a mouse-specific pathogen that shares a virulence strategy and effectors with EPEC and EHEC. In resistant mice (e.g., C57BL/6), CR causes a mild, self-limiting infection, whereas susceptible mice (e.g., C3H/HeN and C57BL/6 *Il22^-/-^*) succumb to the infection^9–14^. The CR infection cycle in C57BL/6 mice progresses through four distinct phases: Establishment (1–3 days post-infection [dpi]) - a small proportion of the inoculum colonizes the cecal lymphoid patch. Expansion (4–8 dpi) - CR colonizes and rapidly expands in the distal colon; first signs of histopathological mucosal damage (including disruption of tight junctions) appear, leading to colonic crypt hyperplasia (CCH), group 3 innate lymphoid cells (ILC3) produce IL-22 and IL-22-regulated antimicrobial proteins (e.g. Reg3γ, Reg3β, LCN2) are detected^15^. Steady-state (8–13 dpi) - shedding plateaus at approximately 10⁹ colony-forming units (CFUs) per gram of feces (GoF) and IL-17A and IL-22 are secreted from neutrophils and Th17/Th22 cells^14,16,17^. In the absence of IL-22 (i.e., infection of *Il22^-/-^* mice) CR-infected mice succumb^14,18^. Clearance (14–20 dpi) - CD4^+^ T cells transition from secretion of IL-17A/IL-22 to predominant IFN-γ production^19,20^, a shift that appears beneficial to the host as *Ifng*^−/−^ mice exhibit delayed pathogen clearance^21^, CR is cleared via IgG-mediated opsonization and neutrophil-driven phagocytosis, and subsequent bacterial out competition by commensal microbiota^22,23^. Infection of C3H/HeN mice follow an accelerated trajectory, progressing rapidly through the initial three phases but failing to enter the clearance phase. Instead, these mice succumb to infection around 10 dpi^10^, underscoring the critical role of host factors in determining infection outcomes^24^.

CR encodes 31 T3SS effectors which induce actin polymerization and changes to the cytoskeleton, subvert immune signaling, and disrupt epithelial barrier integrity^25^. While early studies focused on characterizing individual T3SS effectors, they often overlooked the broader network-level interactions that allow pathogens to retain virulence despite effector deletions. Using a systematic sequential effector deletion programme, we have recently showed that T3SS effectors function as interdependent networks^26^. The effector network concept has later been expanded to *S*. Typhimurium^27,28^. In CR, Tir, EspZ, and NleA are essential effectors, as deletion of any one of them individually rendered CR non-infectious^26^. However, a mutant strain encoding an effector network lacking 19 of the 31 effector genes (CR_14_) maintained colonization, with bacterial shedding above our set threshold of 10^7^ CFU/GoF^26^. Similarly, a strain lacking 10 effectors involved in immune subversion (NleB, NleC, NleE, NleD1/2, NleF, NleH, EspJ, EspL, and EspT), designated CR_i9_, was also shed above the set threshold. To describe scenarios where an effector becomes essential only in a specific perturbed network, we introduced the concept of context-dependent effector essentiality (CDEE)^26^.

Despite significant progress in understanding T3SS effectors *in vivo*, key questions remain regarding their functional network interactions. It is still unclear what is the role of the accessory effectors, and why seemingly nonessential effectors are evolutionarily maintained. Here, using multiple minimal effector networks, we define a subset of 12 accessory CR effectors that are dispensable for colonization yet shape infection outcomes. By generating a strain lacking these 12 effectors (CR_M12_), we reveal their role in modulating immune responses, epithelial barrier integrity, and disease severity across host backgrounds. These findings suggest that maintenance of accessory effectors within a network is shaped by host-driven evolutionary pressure and highlight the presence of a core effector network essential for pathogenesis. Our study advances the understanding of T3SS effector network flexibility, revealing how accessory effectors shape infection dynamics. These findings provide broader insights into bacterial adaptation and host-pathogen interactions.

## Results

### Identification of the accessory CR effectors

To determine the makeup of the accessory effectors within the CR effector network, we generated additional mutants by sequentially deleting effector genes (Fig. 1A), which were tested for their ability to colonize C57BL/6 mice (Fig. 1B). Using our set shedding threshold of 10⁷ CFUs/GoF, we generated the effector network CR_i17_, (an extension of the previously reported CR ^26^) missing 18 effectors, which reached the robustness limit (i.e. deletion of any of the remaining effector genes resulted in shedding below the threshold). We also generated the network CR_P20_ lacking 20 effectors, which also reached the robustness limit (Fig. 1C-E). While a triple *nleG* mutant (CRΔ*nleG1*/*7*/*8*) colonized C57BL/6 mice robustly (Fig. S1A), NleG8, but not NleG1 or NleG7, displayed CDEE in the CR_i_ and CR_P_ network intermediates (Fig. 1C-E).

**Figure 1.**
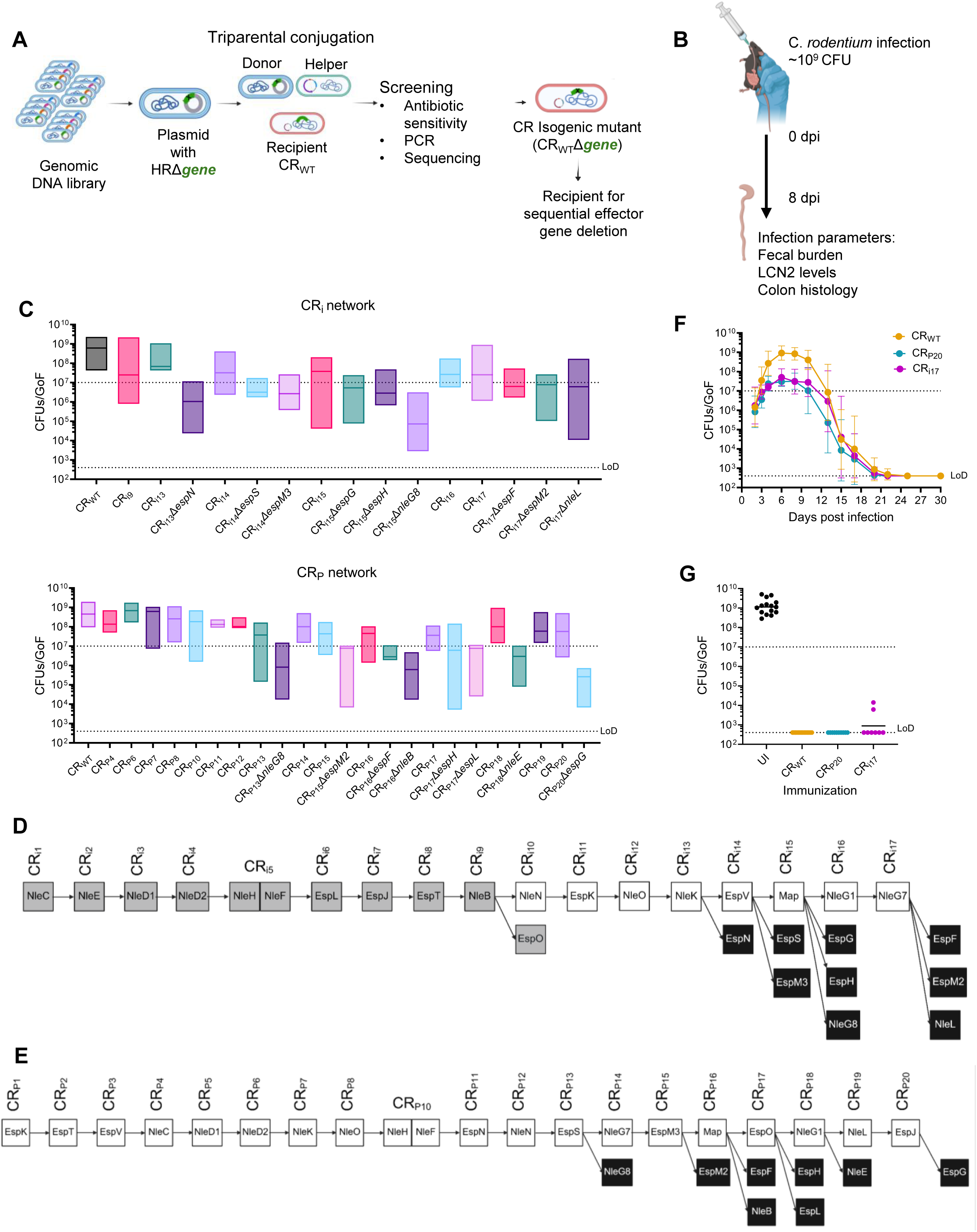
Generation of minimal effector networks. (**A**) A schematic illustrating construction of the effector networks using sequential effector deletion. (**B**) A schematic depicting infection of C57BL/6 mice with CR_WT_ or isogenic mutants. At 8 dpi, infection outcomes were assessed by quantification of CFUs/GoF, fecal LCN2 and CCH. (**C**) CFUs/GoF of the CR_i17_ and CR_P20_ intermediates. Results show median from biological replicates (n ≥ 4 mice per group). (**D** and **E**) Pictorial representations of the effectors deleted in each round of sequential deletion; gray boxes represent CR ^27^, white and black boxes represent that network mutant shedding above or below the threshold, respectively. (**F**) Temporal fecal bacterial shedding in mice infected with CR_WT_, CR_P20_, and CR_i17_. Lines represent the mean bacterial load with each point representing geometric mean ± geometric S.D. from biological replicates (N ≥ 2). (**G**) Fecal CR_WT_ shedding 8 days after reinfection of mice pre-infected with CR_WT_, CR_i17_ or CR_P20_. Shown are geometric mean of biological replicates (N ≥ 2). Each data point represents a single mouse. In **C**, **F**, and **G**, limit of detection (LoD) and the colonization threshold are indicated by dotted black lines. Refer to Table S2 for the exact number of mice used for experimental groups.

CR_i17_ and CR_P20_ retained 13 and 11 of the 31 CR effectors, respectively. A direct correlation between fecal shedding, fecal LCN2 levels and CCH was observed following infection with the CR_i17_ and CR_P20_ intermediates (Fig. S1B-C). CR_i17_ and CR_P20_ transitioned through the four infection phases similar to CR_WT_, with peak shedding at 8 dpi, rapid clearance from 12 dpi, and infection resolution by 22 dpi (Fig. 1F). Moreover, mice infected with either CR_i17_ or CR_P20_ were protected from a secondary challenge with CR_WT_, suggesting that infection with CR_i17_ and CR_P20_ triggered protective immunity (Fig. 1G).

Comparing the effector network compositions of CR_i17_, CR_P20_, and CR_14_^26^ revealed the absence of 12 shared effectors —Map, NleD1/NleD2, NleH, NleF, EspJ, NleG1, NleG7, EspV, EspK, NleN, and NleK— (Fig. 2A). The functions and interacting partners of these effectors are summarized in Table S1 and Figure S1D. To determine the collective impact of these accessory effectors, we generated CR_M12_, a strain lacking the 12 effector genes (Fig. 2B). CR_M12_ maintained robust colonization, which followed a similar infection trajectory as CR_WT_ (Fig. 2C), and displayed protection against secondary challenge with CR_WT_ (Fig. 2D).

**Figure 2.**
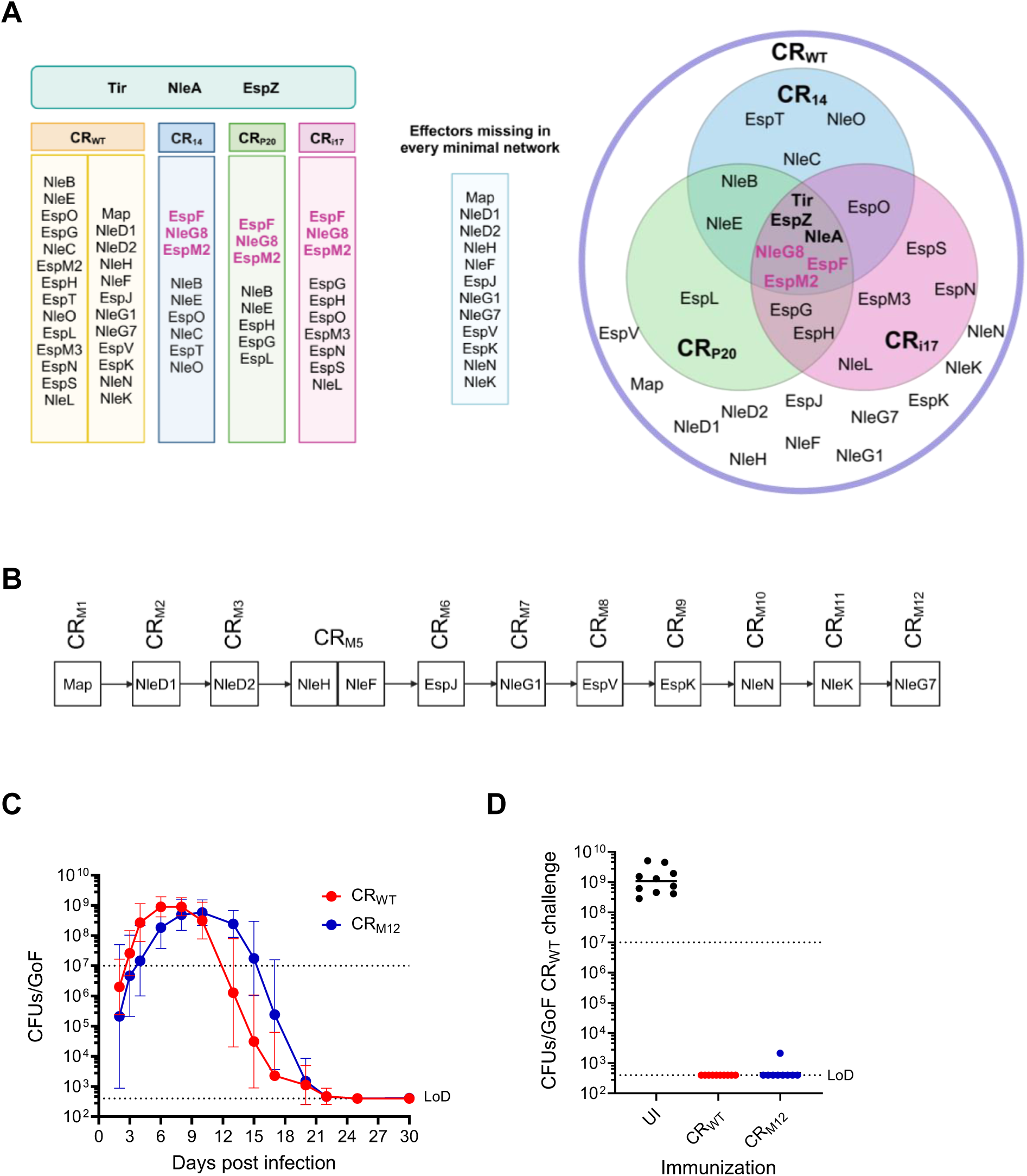
Identification of the accessory effectors. (**A**) Table (**left**) and Venn diagram (**right**) representing effectors present in the CR_WT_, CR_14_, CR_P20_, and CR_i17_ networks. The essential effectors are shown at the top of the table and center of the Venn diagram. The common effectors EspF, NleG8 and EspM2 are shown in pink. Effectors missing from CR_14_, CR_P20_, and CR_i17_ networks are listed in the light blue box. **(B)** Pictorial representation of the rounds of sequential deletion towards the generation of CR_M12_. (**C**) Temporal fecal bacterial shedding of mice infected with CR_WT_ or CR_M12_. Lines represent the mean bacterial load with each point representing geometric mean ± geometric S.D. from biological replicates (N = 3). (**D**) Fecal CR_WT_ shedding 8 days after reinfection of mice pre-infected with CR_WT_ or CR_M12_. Shown are geometric mean of biological replicates (N ≥ 2). Each data point represents a single mouse. In **C** and **D**, LoD and the colonization threshold are indicated by dotted black lines. Refer to Table S2 for the exact number of mice used for experimental groups.

### CR_M12_ is virulent in a susceptible mouse strain

Given that the accessory effectors are dispensable for colonization in C57BL/6 mice, we next assessed if CR_M12_ remains virulent in C3H/HeN mice, where CR infection induces severe weight loss, watery diarrhea, dehydration, and colonic barrier damage^10,24,29^. While 100% of CR_WT_-infected C3H/HeN mice reached the humane endpoint by 10 dpi, CR_M12_ infection resulted in predicted mortality of 83% (Fig. 3A-B). Both CR_WT_ and CR_M12_ exhibited comparable bacterial shedding (Fig. 3C) and induced similar increases in fecal water content compared to uninfected (UI) mice (Fig. 3D). Necropsy at the humane endpoint revealed that infection with both CR_WT_ and CR_M12_ induced severe colonic inflammation, characterized by colon shortening, thickening, and an increased weight-to-length ratio (Fig. 3E-G). Histological analysis of CR_M12_- infected colons revealed extensive mucosal hyperplasia, dense immune cell infiltration, and submucosal thickening, consistent with colonic pathology observed in CR_WT_-infected C3H/HeN mice (Fig. 3H-I). These findings show that despite lacking the 12 accessory effectors, CR_M12_ retains pathogenic potential in a susceptible mouse strain, confirming their accessory ascription.

**Figure 3.**
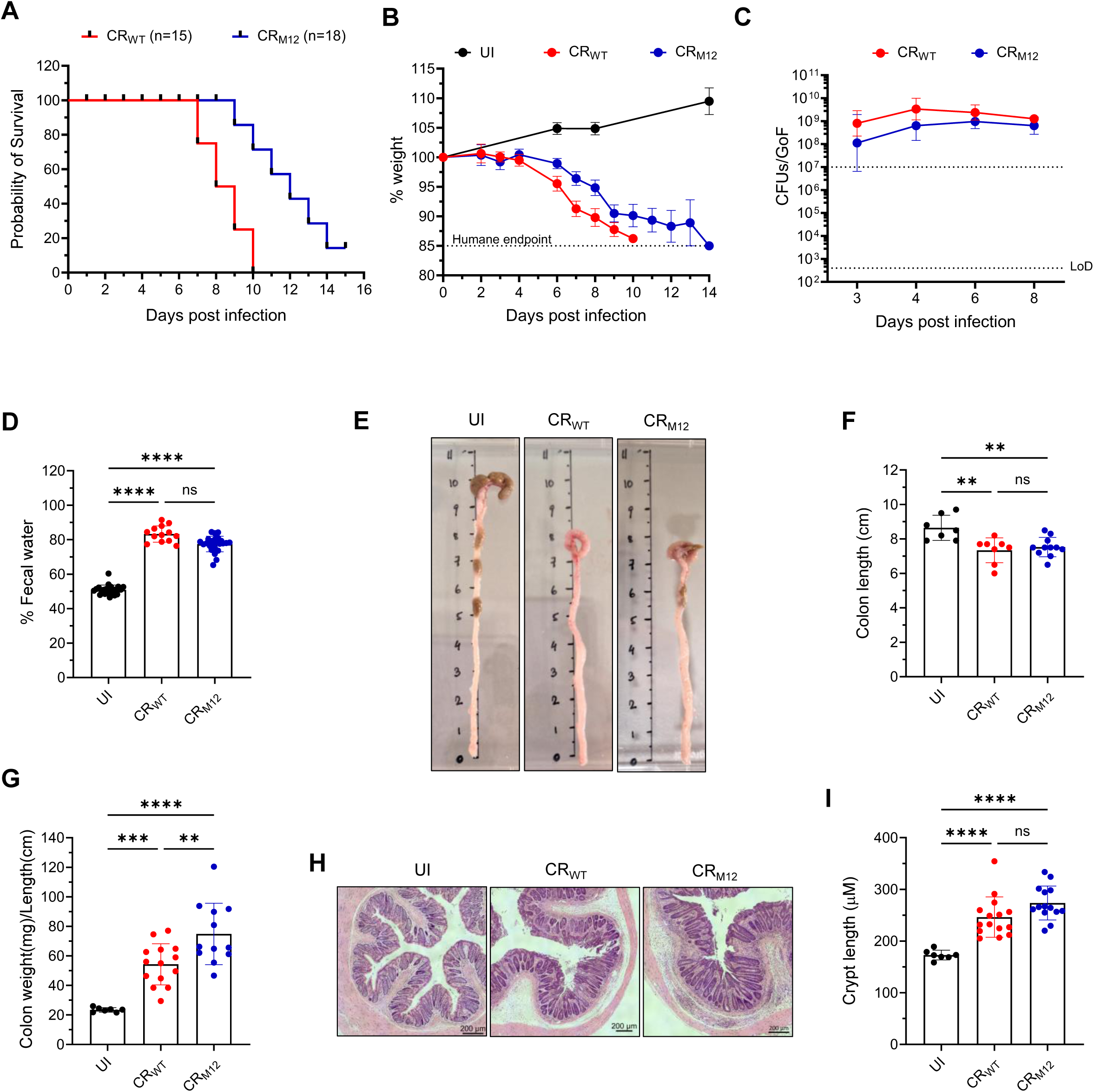
CR_M12_ is virulent in C3H/HeN mice. C3H/HeN mice were infected with either CR_WT_ or CR_M12_. (**A**) Probability of survival and (**B**) temporal weight loss of mice infected with the indicated strains and UI control. Data represents mean ± SEM from biological replicates (N = 3). (**C**) Temporal fecal bacterial shedding. Results show geometric mean ± geometric S.D. from biological replicates (N ≥ 3). The colonization threshold and LoD CFUs/GoF are indicated by dotted lines. (**D**) Fecal water content at 8 dpi (N=3). (**E**) Representative images of distal colons from mice harvested at humane endpoint and UI controls (N ≥ 2). (**F**) Colon length. (**G**) Colon weight-to-length ratio. **(H)** Representative hematoxylin and eosin (H&E)-stained colon sections, and **(I)** crypt-length measurements (scale bars, 200 μm). For **D**, **F**, **G** and **I**, each data point represents a single mouse, and results show mean ± S.D. from biological replicates (N ≥ 2). Statistical significance was determined by one-way ANOVA with Tukey’s multiple comparison test. **p < 0.01; ***p < 0.001; ****p < 0.0001; ns, not significant. Refer to Table S2 for the exact number of mice used for experimental groups.

### CR_M12_ induces lower inflammation in C57BL/6 mice

While CR_M12_ retained its ability to drive severe disease in a susceptible host, we next investigated infection outcome in resistant C57BL/6 mice. At 8 dpi, mucosal-associated bacterial loads were comparable between CR_WT_ and CR_M12_ infected mice (Fig. 4A), and both strains induced similar increases in fecal water content compared to UI mice (Fig. 4B). However, colon weight-to-length ratios were significantly lower in CR_M12_-infected mice compared to CR_WT_-infected controls (Fig. 4C), indicating a reduced inflammatory response^30^. Additionally, CR_M12_ did not induce CCH or expansion of the PCNA-positive zone, indicative of reduced epithelial proliferation^15^ (Fig. 4D-G). In agreement to the role of neutrophils in damage of the intestinal barrier^29^, fecal levels of LCN2/NGAL and S100A8, antimicrobial proteins mainly expressed by neutrophils, were significantly lower following CR_M12_ infection (Fig. 4H-I).

**Figure 4.**
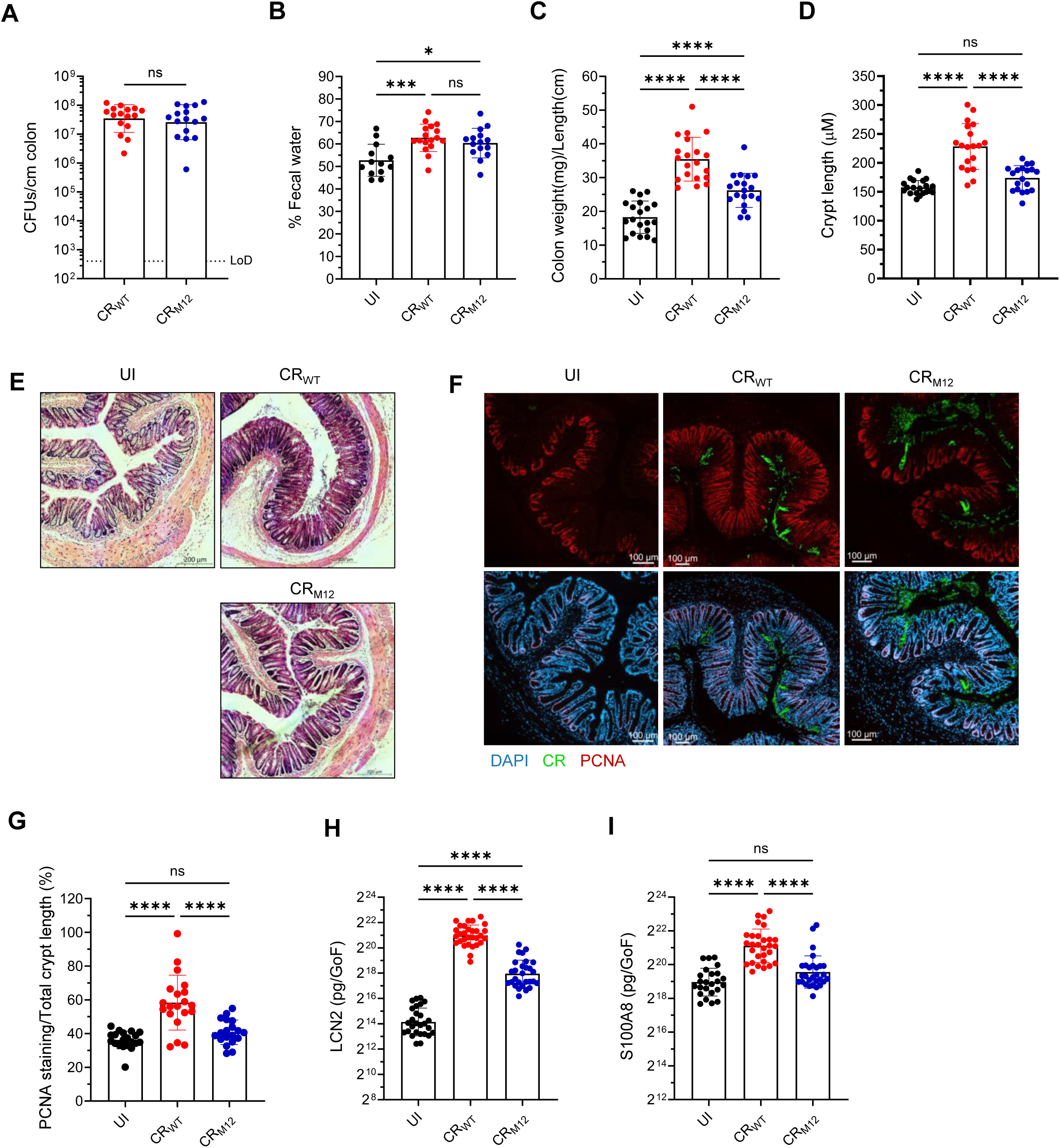
CR_M12_ induces mild tissue damage in C57BL/6 mice. C57BL/6 mice infected with CR_WT_ or CR_M12_, and UI controls, were harvested at 8 dpi. (**A**) Tissue-associated CR (CFUs/cm colon). Results show geometric mean ± geometric S.D. from biological replicates (N = 4). LoD is indicated by dotted black line. (**B**) Fecal water content. (**C**) Colon weight-to-length ratio. (**D**) Crypt length measurements and (**E**) representative H&E-stained colon sections (scale bars, 200 μm) (N=4). In **D**, each dot represents the mean per mouse. **(F)** Representative immunostaining images of colonic sections, and **(G)** quantification of PCNA staining as a percentage of total crypt length (N=4). Each dot represents the mean per mouse; CR (green), DAPI (blue), PCNA (red) (scale bars, 100 μm). Fecal (**H**) LCN2 and (**I**) S100A8 levels. For **A**-**D**, **G**-**I**, each data point represents a single mouse, and results show mean ± S.D. from biological replicates (N = 4). Statistical significance was determined by a two-tailed unpaired t-test for **A**, and one-way ANOVA with Tukey’s multiple comparison test for **B**-**D**, and **G**-**I**. *p < 0.05; ***p < 0.001; ****p < 0.0001; ns, not significant. Refer to Table S2 for the exact number of mice used for experimental groups.

These results suggest that despite similar level of attachment to the colonic mucosa, CR_M12_ infection triggers reduced tissue damage and inflammation.

Compared to CR_WT_, infection with CR_M12_ triggered reduced colonic secretion of Th1 and Th17 cytokines IFN-γ, IL-22, IL-17A, and inflammasome-dependent IL-1β, involved in stimulating IL-22 responses^31^, correlating with the observed lower tissue damage (Fig. 5A-D). However, comparable levels of TNF, IL-6, IL-10, CXCL1, CCL3, GM-CSF, and G-CSF were observed upon CR_WT_ and CR_M12_ infection (Fig. S2A-G). Interestingly, colonic IL-18 levels, an inflammasome-dependent cytokine that can also be released by non-professional immune cells like IECs^32^, were significantly higher in CR_M12_-infected mice compared to CR_WT_-infected mice (Fig. 5E). Therefore, despite maintaining similar colonization dynamics to CR_WT_, CR_M12_ induces significantly lower levels of pro-inflammatory cytokines compared to CR_WT_ infection.

**Figure 5.**
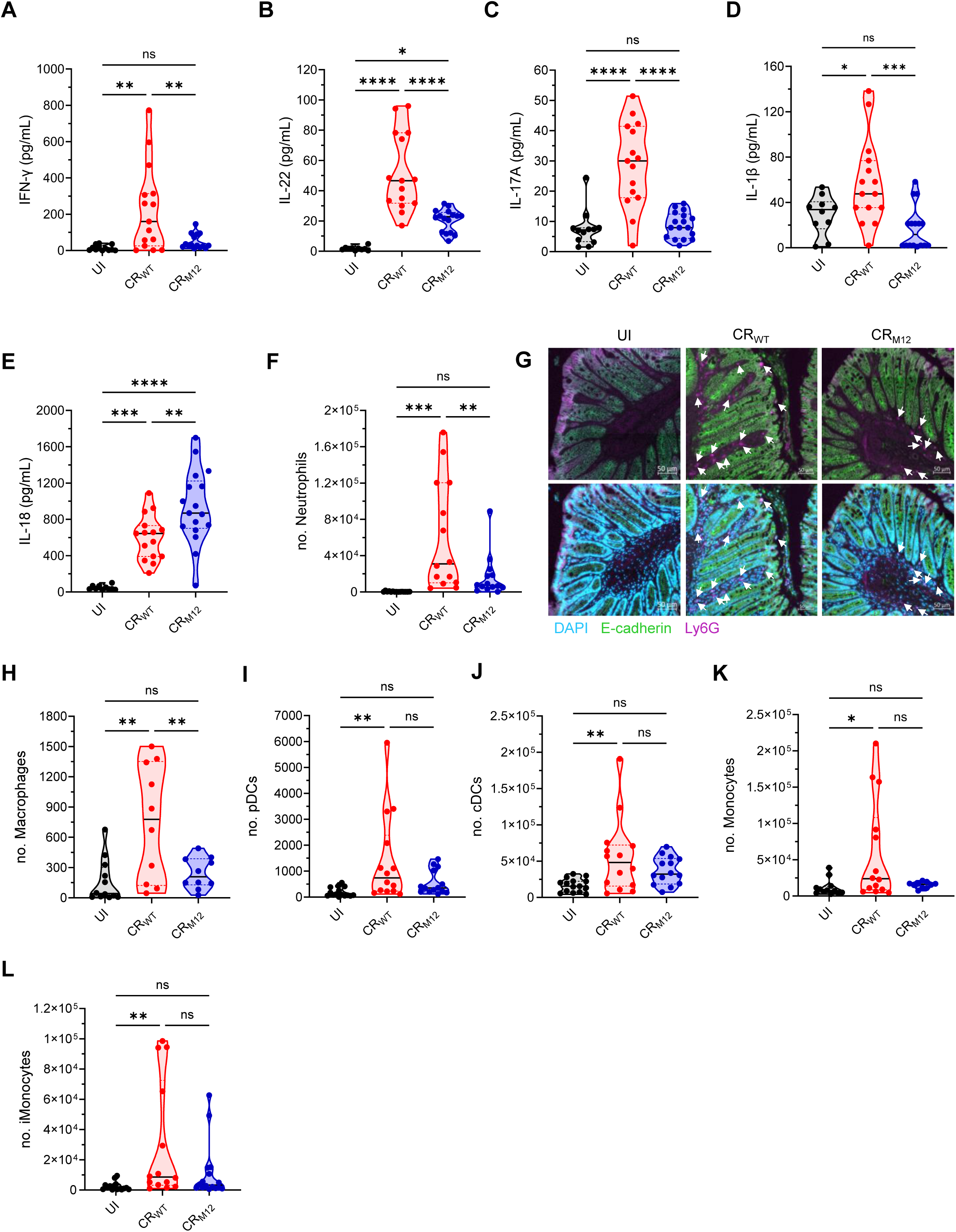
CR_M12_ triggers lower inflammation in C57BL/6 mice. C57BL/6 mice infected with CR_WT_, or CR_M12_, and UI controls were harvested at 8 dpi. Mucosal secretion of (**A**-**E**) IFN-γ, IL-22, IL-17A, IL-1β, and IL-18 cytokines in colonic explant cultures. Results show mean ± S.D. from biological replicates (N = 4). Total numbers of (**F**) neutrophils, (**H**) macrophages, (**I**) pDCs, (**J**) cDCs, (**K**) monocytes, and (**L**) iMonocytes per 200 μl of colon homogenate. Results show mean ± S.D. from biological replicates (N = 3). (**G**) Representative immunostaining images of colonic sections showing neutrophil influx to the colonic mucosa (N=4). The intestinal epithelial cells express E-cadherin (green) and are DAPI^+^ (blue), whereas neutrophils are Ly6G^hi^ (violet) and DAPI^+^, but E-cadherin^-^ (scale bars, 50 μm). The upper panel is a merged image of E-cadherin and Ly6G and the lower panel shows merged images of all three channels (DAPI, E-cadherin, and Ly6G). For **A**-**F**, and **H**-**L,** each data point represents a single mouse; statistical significance was determined by one-way ANOVA with Tukey’s multiple comparison test. *p < 0.05; **p < 0.01; ***p < 0.001; ****p < 0.0001; ns, not significant. Refer to Table S2 for the exact number of mice used for experimental groups.

Since CR_M12_ infection resulted in lower tissue damage and attenuated immune responses, we next examined immune cell recruitment to the colon at the peak of infection. While B cells, CD4^+^, and CD8^+^ T cells were recruited at similar levels (Fig. S3A-F), neutrophil and macrophage infiltration was significantly lower following CR_M12_ infection (Fig. 5F-H, S4A-B). This was validated by Ly6G staining of thin colonic sections, which showed reduced neutrophil accumulation (Fig. 5G). Additionally, CR_M12_-infected colons lacked the expansion of plasmacytoid dendritic cells (pDCs), conventional dendritic cells (cDCs), monocytes, and inflammatory Ly6C^+^ monocytes (iMonocytes) which were observed in CR_WT_ infection (Fig. 5I-L, S5A-C). Together, these findings suggest that while not affecting colonization, the 12 accessory effectors alter infection outcomes, impacting on tissue damage and host immune responses.

### CR_M12_ causes lower epithelial barrier disruption

Given the observed reduction in tissue damage and immune activation in CR_M12_ infected mice, we next determined whether this phenotype was linked to effector-mediated modulation of colonic IECs (cIECs). To this end, we specifically FACS sorted cIECs colonized by CR_M12_ and CR_WT_ (EpCAM^+^/CR^+^) at 6 dpi (Fig. 6A). We selected this time point as the expansion phase is associated with T3SS activity and effector translocation^15^. Deep quantitative proteomic analysis of EpCAM^+^/CR^+^ cIECs was performed (Fig. 6A) and principal component analysis (PCA) revealed distinct clustering between infected and UI mice (PC1, 80.7% variance), with additional separation between CR_WT_ and CR_M12_ infection (PC2, 5.5% variance) (Fig. 6B). We quantified the abundance of 8040 mouse proteins, with hierarchical clustering analysis showing that our biological replicates clustered by treatment (UI vs CR_WT_ vs CRM_12_) (Fig. 6C). We recorded no global differences in protein abundances, with only 126 proteins being different between CR_WT_ and CR_M12_ infection (p < 0.05, |log2FC| > 0.5) (Fig. 6C and 6D). This suggests that CR_M12_ infection largely mirrors CR_WT_ at the cellular level despite lacking ∼39% of its T3SS effector repertoire, emphasizing the robust nature of the effector network.

**Figure 6.**
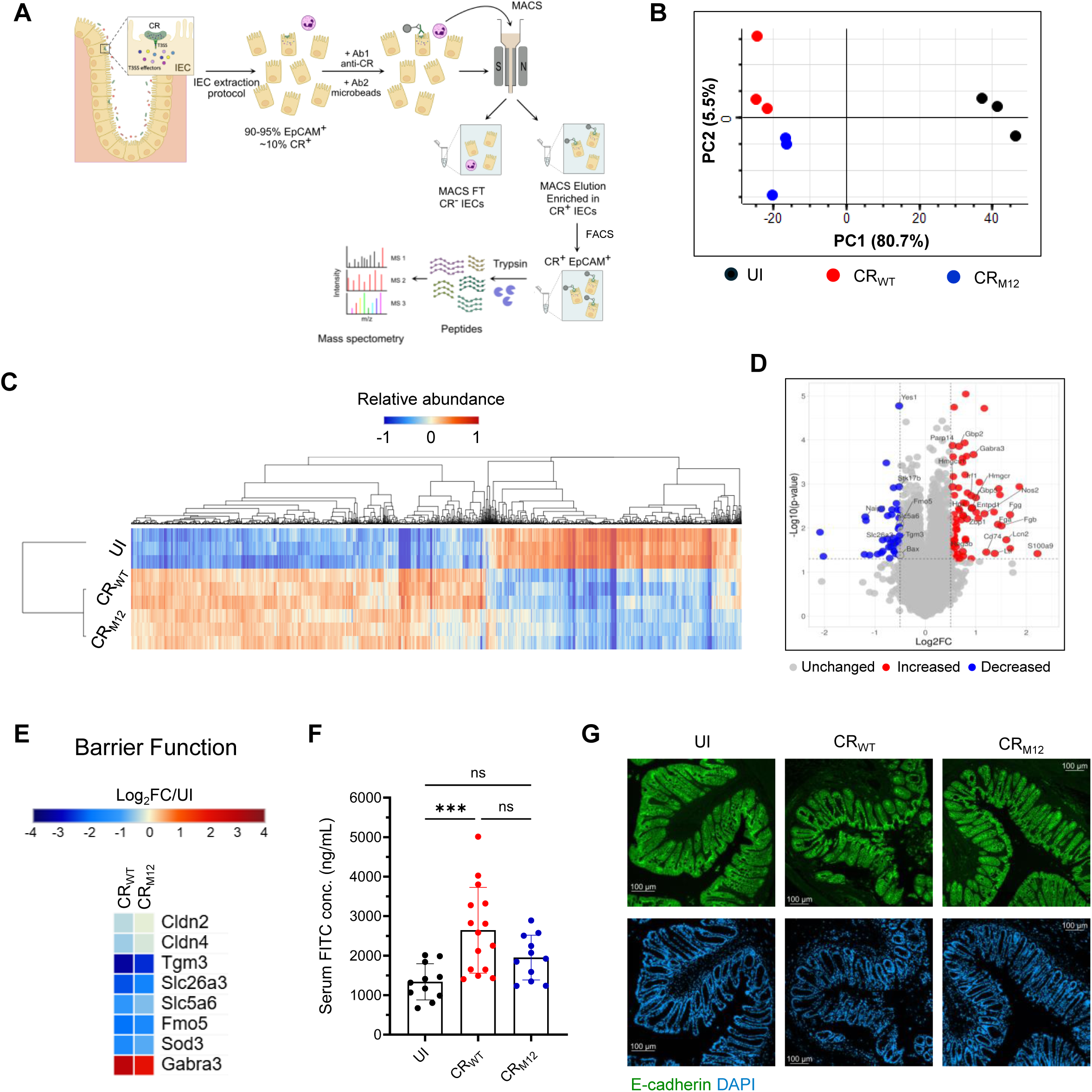
CR_M12_ triggers lower epithelial barrier disruption in infected IECs. (**A**) A schematic illustrating sample preparation leading to proteomics analyses of EpCAM^+^/CR^+^ IECs (**B**) PCA of mouse proteins in EpCAM^+^/CR^+^ isolated from UI mice and mice infected with CR_WT_ and CR_M12_ (N=3). (**C**) Hierarchical clustering (one minus cosine similarity) of relative abundance of 4197 proteins significantly changing with CR_WT_ and/or CR_M12_ infection compared to UI. (**D**) Volcano plot (Log_2_FC of CR_WT_ vs CRM_12_) showing significantly (y-axis), and differentially (x-axis) regulated proteins between CR_WT_ and CR_M12_ infection. (**E**) Heatmap of selected proteins involved in regulation of barrier integrity. **(F)** Intestinal permeability was measured at 8 dpi by determining FITC-dextran levels in the serum. Each data point represents a single mouse, and results show mean ± S.D. from biological replicates (N = 4). Statistical significance was determined by one-way ANOVA with Tukey’s multiple comparison test. ***p < 0.001; ns, not significant. **(G)** Representative immunostaining images of colonic sections from N= 4 biological repeats showing epithelial barrier disruption. The intestinal epithelial cells express E-cadherin (green) and are DAPI^+^ (blue), (scale bars, 100 μm). The upper and lower panel show E-cadherin and DAPI staining, respectively. Refer to Table S2 for the exact number of mice used for experimental groups.

We focused our pathway analysis on processes that might explain the observed differences in infection outcomes. This revealed that both CR_WT_ and CR_M12_ inhibited cIEC cell death (apoptotic, pyroptotic and necroptotic) (Fig. S6A). While we detected higher abundance of Fas, the levels of the downstream death adaptor proteins TRADD and FADD, targeted by NleB^33^ were reduced in cIECs infected with either CR_WT_ or CR_M12_. In contrast, the abundance of the anti-apoptotic factors BIRC5 and HSP90B was enriched. Consistent with the reduced immune cell infiltration and cytokine secretion, CR_M12_-infected IECs exhibited lower expression of cytokine-regulated pro-inflammatory and antimicrobial proteins (Fig. S6B). In particular, lower expression of IL-22-regulated proteins S100A9, LCN2, REG3β, SAA4 and IFN-γ- regulated CD74^34^, GBP2 and GBP5^35^ was observed in CR_M12_- compared to CR_WT_-infected IECs (Fig. 6D, Fig. S6B).

Proteomics analysis was indicative of CR_M12_ infection resulting in a comparatively milder disruption of barrier function than CR_WT_ (Fig. 6E). Particularly, CR_WT_ infection was characterized by loss of key structural proteins—CLDN2, CLDN4^36^ and their regulators such as TGM3, & FMO5 involved in mucus barrier formation and stabilization^37,38^; SLC26A3, SLC5A6, and SOD3 involved in maintaining the integrity and stability of epithelial tight junction barrier^39–43^ and an increase in barrier-disrupting factor: GABRA3^44^. The changes in these markers observed in CR_M12_-infected cIECs were less dramatic, suggesting reduced barrier disruption (Fig. 6E). To validate these results, we performed the FITC-dextran intestinal permeability assay^45^. This revealed that infection with CR_M12_ resulted in significantly lower levels of FITC-dextran permeating into the serum compared to those in CR_WT_-infected mice; moreover, immunofluorescence staining for E-cadherin revealed lower epithelial damage (Fig. 6F-G). These results suggest that infection of CR_M12_ causes less disruption of the cIEC monolayer, which in turn may result in lower IL-22 responses.

### CR_M12_ is virulent in mouse model of impaired intestinal barrier repair

Since the level of barrier disruption and IL-22 differentiates infection with CR_WT_ and CR_M12_, we investigated infection outcomes in C57BL/6 *Il22^-/-^*mice, which have impaired barrier repair mechanisms and succumb to CR_WT_ infection^13,14^. Importantly, *Il22^-/-^* mice survive infection with a CR strain lacking EspF, which does not disrupt the epithelial barrier^13^. We therefore investigated how the barrier disruption caused by CR_M12_ impacts disease progression in *Il22^-/-^* mice.

While 100% of CR_WT_-infected *Il22^-/-^* mice succumbed to infection, CR_M12_ infection resulted in a predicted mortality of 67% (Fig. 7A-B). However, both CR_WT_ and CR_M12_ exhibited comparable bacterial shedding and induced similar increases in fecal water content (Fig. 7C-D). Necropsy at the humane endpoint revealed severe colonic inflammation, substantial shortening and thickening of the colon, and an increased colon weight-to-length ratio, indicating that CR_M12_- infected mice that succumb to infection likely suffer from the same unresolved damage to the barrier as CR_WT_-infected mice (Fig. 7E-G). Consistently, histological analysis showed extensive CCH, large immune cell infiltration, and submucosal thickening, confirming severe intestinal pathology (Fig. 7H-I). Moreover, CR_WT_ and CR_M12_ infections resulted in comparable systemic dissemination to liver and spleen (Fig. S7A-B). Likewise, both infections triggered splenic inflammation and spleen weight gain, further indicating systemic immune activation (Fig. S7C-D). These findings suggest that CR_M12_ induces disease and inflammation similar to CR_WT_ when the ability to restore the barrier integrity is compromised.

**Figure 7.**
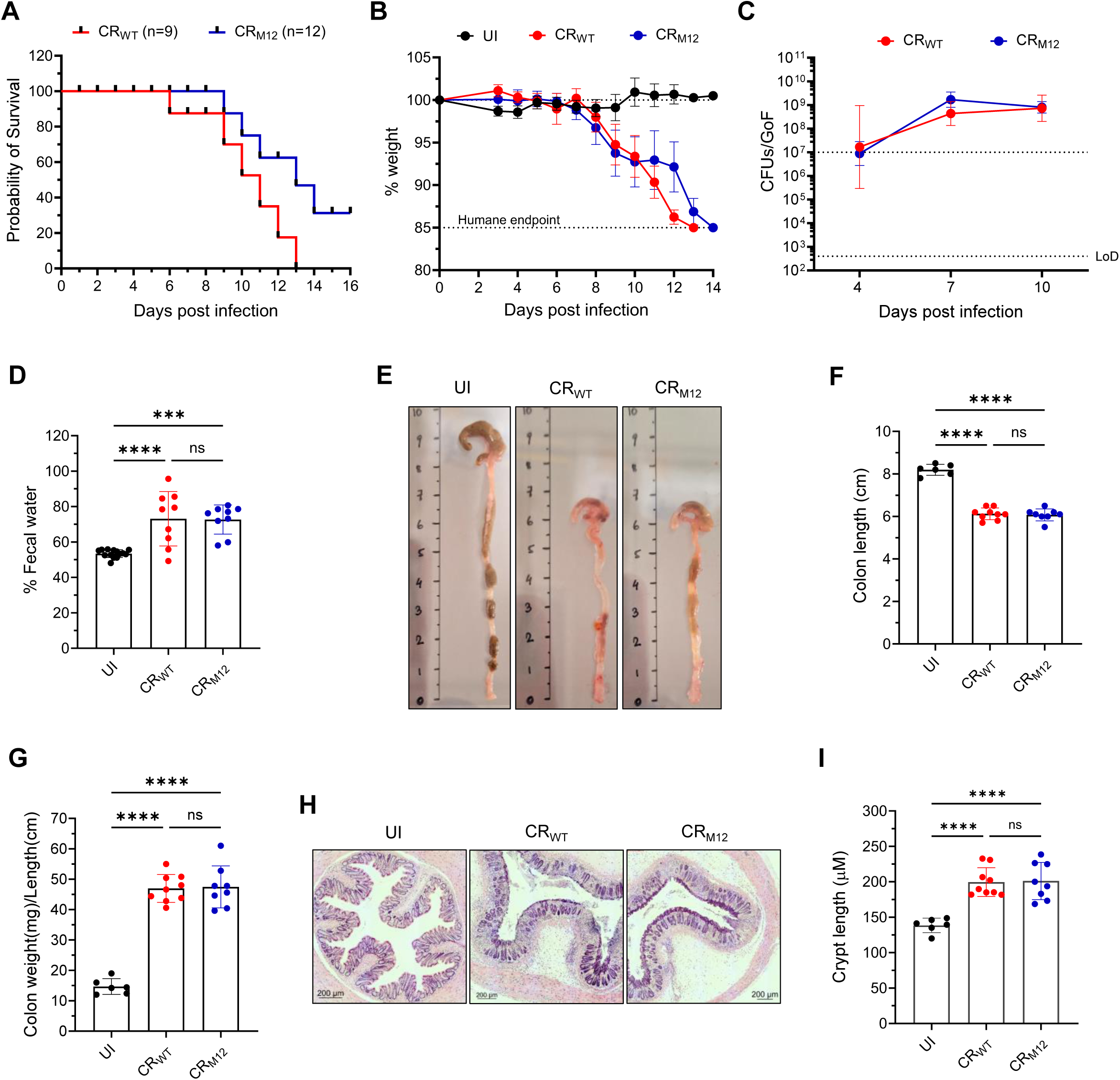
CR_M12_ infection results in mortality of mice with impaired barrier repair. C57BL/6 *Il22^-/-^* mice were infected with either CR_WT_ or CR_M12_. (**A**) Probability of survival and (**B**) temporal weight loss of infected mice and UI control. Data represents mean ± SEM from biological replicates (N = 2). (**C**) Temporal fecal bacterial shedding. Results show geometric mean ± geometric S.D. from biological replicates (N = 2). The colonization threshold and LoD are indicated by dotted black lines. (**D**) Fecal water content at 7 dpi. (**E**) Representative images of distal colons from mice harvested at humane endpoint and of UI controls (N = 2). (**F**) Colon length. (**G**) Colon weight-to-length ratio. (**H**) Representative H&E-stained colon sections, and (**I**) crypt-length measurements (scale bars, 200 μm). Each dot represents the mean per mouse. For **D**, **F**, **G** and **I**, each data point represents a single mouse, and results show mean ± S.D. from biological replicates (N ≥ 2). Statistical significance was determined by one-way ANOVA with Tukey’s multiple comparison test. ***p < 0.001; ****p < 0.0001; ns, not significant. Refer to Table S2 for the exact number of mice used for experimental groups.

## Discussion

Defining the minimal set of genes required for bacterial survival and pathogenesis has been a central theme in bacterial evolution and pathogen-host interactions^27,28,46–52^. In this study, we further demonstrated the flexibility of T3SS effector networks, which challenge the traditional view of a fixed set of essential virulence factors. Instead, our data highlights a modular, adaptable effector network where different effectors assume context-dependent roles.

By generating the minimal effector networks CR_P20_, CR_i17_ and CR_14_^26^, we uncovered that NleG8 exhibits CDEE. NleG8’s essentiality in specific effector networks may stem from its interaction with GOPC, a regulator of tight junction integrity and mucosal homeostasis, enabling it to modulate host signaling pathways^53^. While dispensable in CR_WT_, NleG8 became critical in CR_i17_ and CR_P20_ intermediates, illustrating the compensatory dynamics within the effector network. Moreover, EspF, which exhibited CDEE in the CR_14_ network^26^, also showed essentiality during the construction of CR_P20_ and CR_i17_ network. EspF, like the 12-effector network absent in CR_M12_, plays a pivotal role in disrupting tight junctions, cytoskeletal remodeling, and immune evasion^54^. Taken together, these data show that disruption of tight junctions during infection with A/E pathogens is a key trigger of inflammatory responses. Yet, compared to deletion of *espF* alone^13^, deleting the 12 accessory effectors has a milder effect on infection outcomes as was illustrated by infection of *Il22^-/-^*mice, highlighting the potential hierarchy of effectors in terms of essentiality during infection.

By systematically identifying a subset of effectors dispensable for infection, this study takes a step towards defining the core effector network required for pathogenesis. While the accessory effectors are not essential for colonization, their deletion significantly altered immune responses, and epithelial integrity. The evolutionary persistence of accessory effectors such as Map, NleD, EspJ, NleH, and NleF, also typically found in EPEC and EHEC isolates^8,26,55^, suggests that they provide a selective advantage in optimizing pathogen-host interactions, rather than serving as redundant elements. Proteomic profiling of infected cIECs revealed significant shifts in barrier integrity regulators, which we validated functionally. The increase in IL-18 secretion observed in CR_M12_-infected mice likely results from the deletion of NleF, which normally suppresses this cytokine’s processing and secretion^56^. This increase combined with the lower tissue damage could make CR_M12_ a good tool to study the effect of local IL-18 responses in colitis, given its role in intestinal tolerance^57^. Similarly, the absence of Map in CR_M12_ may contribute to the observed milder disruption of gut barrier function^58^.

Despite CR_M12_’s attenuated immune activation in resistant hosts, it retained pathogenic potential in susceptible models of infection, albeit with lower mortality rates likely ensuing from its reduced tissue damage capability. This adaptability, illustrated by the fact that the remaining 61% of effectors in CR_M12_ are sufficient to trigger disease, mirrors findings in other pathogens, such as *Salmonella* Typhimurium, where cooperation between effector genes within a network govern tissue tropism^27^.

Effector network plasticity is a key determinant of host adaptation, enabling pathogens to exploit novel immune landscapes and establish infection in new hosts^8,26,27,59,60^. The variability of T3SS effector repertoires across pathogenic bacteria further illustrates the adaptability of these systems. Human pathogens such as EPEC and EHEC share homologous effectors with CR yet exhibit differences in effector composition, reflecting host- and tissue-specific adaptations. For instance, while effectors like Tir and EspF are conserved and critical for host colonization, others are variably present or exhibit mutations that modulate their functionality^55,61^. Moreover, clinical isolates of EPEC and EHEC often lack certain effectors or carry non-functional alleles yet retain pathogenic potential^8^. Notably, NleG, EspM and EspJ, are more commonly found in EPEC and EHEC isolates that are linked to severe human infections^6,8^.

In conclusion, our study highlights bacterial pathogenesis as a highly adaptable, host-responsive process shaped by a flexible effector network. By identifying a subset of 12 accessory effectors, we refine the framework for distinguishing essential from accessory virulence determinants, establishing a foundation for future research aimed at fully defining the core effector network. Understanding the selective pressures governing effector composition variability may provide insights into host adaptation, pathogen evolution, and effector-driven immune evasion. These findings also have broad implications for developing targeted therapeutics, including effector-based interventions that disrupt virulence pathways at critical regulatory nodes.

## Materials & Methods

### Strains

The bacterial strains employed in this study are listed in Table S3. The CR strains were grown at 37 °C in Lysogeny broth (LB), or LB agar plates (15% v/v). Nalidixic acid (Nal, 50 μg/ml), streptomycin (Sm, 50 μg/ml), gentamicin (10 μg/ml), kanamycin (Kan, 20 μg/ml) were added for plasmid or strain selection, as required.

### Generation of mutants

*Escherichia coli* CC118-λpir harboring pSEVA612S recombinant plasmid was used for propagation of plasmids. Specific deletion of effector genes via homologous recombination was performed as described in Ruano-Gallego *et al.*^26^. Briefly, 300 base pairs (bp) upstream and downstream flanking region (henceforth named HRΔ*gene*) of the target gene was cloned into pSEVA612S and *E. coli* CC118-λpir was electroporated with the recombinant plasmid pSEVA612S-HRΔ*gene* to generate the donor strain for conjugation. Genes were deleted via tri-parental conjugation, where the helper strain (*E. coli* 1047 pRK2013) was incubated with specific donor strain (*E. coli* CC118-λpir-pSEVA612S HRΔ*gene*) on LB agar at 37 °C, followed by incubation with recipient strain (CR strain carrying pACBSR). Conjugants were selected on LB + Gm + Sm plate. To remove the Gm resistance and for enhanced homologous recombination required for gene deletion, conjugants were grown in LB + Sm + 0.4% L-arabinose for 6-8 h to induce the expression of I-SceI endonuclease from pACBSR and streaked on LB + Sm plates. To remove pACBSR, strains were passaged several times in LB followed by selection of Sm-sensitive bacteria. Deletion mutants were screened by PCR for confirmation of the deletion using Taq 2X Master Mix (NEB) and primers listed in Table S3. All deletion mutants were confirmed by sequencing (Eurofin) (Fig. 1A).

### Mouse experiments

All animal experiments complied with the Animals Scientific Procedures Act 1986 and U.K. Home Office guidelines and were approved by the local ethical review committee. Experiments were designed in agreement with the ARRIVE guidelines^62^ for the reporting and execution of animal experiments, including sample randomization and blinding. Mouse experiments were conducted with five mice per group. Pathogen-free female 18-20g C57BL/6 mice and 8–10- week-old C3H/HeN mice were purchased from Charles River Laboratories. C57BL/6 *ll22*^-/-^ mice were housed and bred in dedicated animal facilities of Imperial College London. All mice were housed in pathogen-free conditions, at 20-22 °C, 30-40% humidity on 12 h of light/dark cycle in high-efficiency particulate air (HEPA)-filtered cages with sterile corn cob bedding, nesting material, and enhancements (chewing toy, and opaque and transparent cylinders), and were fed with RM1 (E) rodent diet (SDS diet) and water ad libitum. All C57BL/6 *IL22^-/-^* mice were genotyped and tested for the presence of iCre using multiplex PCR as previously described^19^. Primer sequences (5’ to 3’) used: Forward, CAGGCTCTCCTCTCAGTTATCA; Wildtype reverse, TCCTGAAGG CCAAAATAGG; Mutant reverse, CCTCAGGTTCAGCAGGGAAC.

### CR infection

Wild type CR (strain ICC169) or its isogenic mutants were grown overnight in LB supplemented with 50 μg/ml nalidixic acid at 37 °C at 180 rpm, were centrifuged at 3000 x g for 10 min, and resuspended in sterile 1X phosphate buffered saline (PBS). Mice weight were recorded (d0) and they were infected with approximately 3 x 10^9^ CFUs in 200 μl sterile PBS by oral gavage as previously described^63^. For mock infection (UI mice), mice received 200 μl sterile PBS. The inocula CFUs were confirmed by CFU quantification as previously described^63^. Infections were followed by plating mouse stools at 2-3 dpi onwards on LB + Nal plates as previously described^63^. For experiments (Fig. 1F, G; 2C, and D), mice infection kinetics were followed until total clearance of the bacteria, defined as two consecutive CR negative stool samples; mice were then reinfected with CR_WT_ strain ICC169 to follow bacterial shedding for an additional 8 days.

### Fecal water content analysis

To determine the fecal water content, feces were freshly collected in pre-weighed 1.5 mL tubes with punctured cap. The tubes with wet feces were weighed and incubated at 55°C. The tubes were weighed everyday till the weight did not change and were recorded. The wet weight and dry weight of feces was determined by subtracting the weight of the tube and fecal water content was estimated using the following equation:

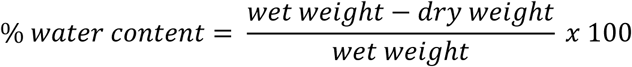

### Postmortem analyses

Mice were monitored for changes in weight every day. Non-susceptible C57BL/6 mice were culled at indicated across various experiments, while susceptible C3H/HeN and immunocompromised *Il22^-/-^* C57BL/6 mice were culled at humane endpoint (Humane end point was met when a mouse lost 15% of its d0 body weight). Postmortem, the large intestine of mice consisting of the caecum and colon was excised, laid on a clean surface in parallel to a mm scale to estimate the colon length, and its picture was taken using a digital camera. Colon was removed from caecum, feces removed and weighed using a digital scale. The weight of the colon was normalized to its length and recorded. From the distal side of the colon, 0.5 cm was stored in 4% paraformaldehyde (PFA) used for histological analysis, next 0.5 cm was used for colon explant (described below). For enumeration of tissue-associated bacteria in colons of infected mice, distal colon was excised, longitudinally cut, feces removed with care without affecting the mucus layer, homogenized in sterile PBS using gentleMACS dissociator (Miltenyi Biotec.), and plated on LB + Nal plates. Similarly, to estimate systemic dissemination of CR, the liver and spleen was harvested, weighed, homogenized in sterile PBS, and plated on LB + Nal plates.

### Histological analysis and immunostaining

Colonic samples were fixed in 4% PFA for 150 min, followed by immersion in 70% ethanol. The fixed tissues were embedded in paraffin, and sectioned at 5 μm. The sections were then stained either with hematoxylin and eosin (H&E) or processed for immunofluorescence. Crypt hyperplasia was quantified in each mouse by measuring >12 well-oriented crypts. The histological sections were blindly evaluated, and the mean crypt length from each mouse was plotted. For immunofluorescence, sections were dewaxed by immersion in Histoclear solution for 10 min x2, followed by immersion in 100% ethanol, for 10 min x2, 95% ethanol for 3 min x2, 80% ethanol for 3 min, and 1X PBS-0.1% Tween-20-0.1% saponin (PBS-TS), for 3 min x2. The sections were heated for 30 min in 0.3% trisodium citrate-0.05% Tween-20 in distilled H_2_O (demasking solution). The slides were washed in PBS-TS, followed by blocking in PBS-TS supplemented with 10% normal donkey serum (NDS) for 20 min. The slides were incubated overnight at 4 °C with anti-CR rabbit polyclonal antibody (1:50), mouse anti-PCNA antibody (1:500), anti- E-cadherin (1:50), and/or anti-Ly6G (1:200) antibody. The following day the slides were washed in PBS-TS, 10 min x2, and incubated with the appropriate secondary antibody (1:100) or DAPI (1:1000) to stain DNA (Table S4). The slides were washed and mounted with ProLong Gold antifade mountant. The sections were imaged and analyzed on Zen 2.3 Blue Version (Carl Zeiss Microimaging GmbH, Germany). PCNA measurements were obtained from >10 well-oriented crypts per mouse. PCNA staining was represented as a percentage of the respective crypt length. Colon sections with less than 10 well-oriented crypts observed were excluded from the analysis.

### Extraction of CR-attached/infected cIECs for proteomics

Female, 8-10 weeks old, C57BL/6 were infected with either CR_WT_ or CR_M12_ (∼3*10^9^ CFUs). UI mice of same age were used as control. The experiment was performed in biological triplicates. In each experiment, at 6 dpi, cIECs were extracted from five or more well-colonized mice or UI controls, pooled and the further processed. Briefly, after necropsy 2 cm distal colon was excised, sliced open longitudinally, feces removed, placed in 4 ml enterocyte dissociation buffer (1X HBSS without Mg and Ca with 10 mM HEPES, 1 mM EDTA, and 0.5% of β- mercaptoethanol) and incubated at 37 °C with shaking at 180 rpm for 45 min. The remaining tissue was removed. Dissociated cIECs were collected by centrifugation (3000 rpm for 10 min, at RT), washed once with ice-cold 1X PBS (2000 x g for 10 min at 4 °C) and resuspended in Mg and Ca-free Dulbecco’s phosphate buffered saline (DPBS) containing 50 µg/ml DNAse I (Thermo Fisher Scientific) and incubated 10 min at room temperature. cIECs were then passed through 70 μM cell strainer to disrupt the tissue, collected by centrifugation, washed with FACS buffer (DPBS supplemented with 5% fetal bovine serum [FBS] and 2 mM EDTA), and incubated in 10% Fc Block (Miltenyi Biotec) in FACS buffer for 10 min on ice. cIECs were stained by anti-CR rabbit polyclonal antibody (at 1:1000 dilution) for 20 min on ice, followed by two washes in MACS (Magnetic activated cell sorting) buffer (DPBS supplemented with 2 mM EDTA and 5% bovine serum albumin [BSA]; filtered). The cells were resuspended in MACS buffer and incubated with anti-rabbit beads for 15-20 min on ice, followed by two washes in MACS buffer and resuspension in the same. LS column fitted to the magnetic field of MACS separator was activated by passing MACS buffer. Cell suspension was then applied into the column followed by three washes with MACS buffer. Magnetically labeled CR^+^ IECs were eluted from the LS column in 5 ml MACS buffer. MACS eluate, enriched in CR^+^ IECs were washed twice in ice-cold PBS, once with FACS buffer, followed by incubation in FACS buffer containing anti-CR rabbit polyclonal antibody (at 1:2000) for 20 min on ice to relabel CR^+^ IECs. Cells were washed and incubated in FACS buffer with anti-rabbit PE and anti- EpCAM-APC antibodies for 30 min on ice. Cells were washed twice in sorting buffer (DPBS without Mg and Ca, supplemented with 1% FBS and 2 mM EDTA) and resuspended in the same. The sample from UI control was used for gating of CR^+^ cells. From MACS eluate, EpCAM^+^/CR^+^ single cells were sorted, (∼[6.1 - 7.9] x 10^5^ for CR_WT_ and [4.6 - 6.9] x 10^5^ for CR_M12_) representing CR-attached/infected IECs. From UI control, ∼10^6^ EpCAM^+^/CR^-^ IECs were sorted which was used as a negative control. The sorted cells were collected by centrifugation and stored at -80 °C until proteomics analysis.

### Sample preparation and TMT labeling

EpCAM^+^/CR^+^ IECs from CR_WT_ or CR_M12_-infected mice and EpCAM^+^/CR^-^ IECs from UI mice from three biologically repeats were solubilized in lysis buffer (100 mM triethylammonium bicarbonate [TEAB], 1% sodium deoxycholate (SDC), 10% isopropanol, 50 mM NaCl) supplemented with Protease and Phosphatase inhibitor cocktail (Thermo Fisher Scientific), boiled for 5 min, and re-sonicated. Protein concentration was determined with Quick Start™ Bradford protein assay (BioRad) according to the manufacturer’s protocol. 15 μg of protein were reduced in 5 mM Tris 2-carboxyethyl phosphine (TCEP) for 1 h, followed by alkylation with 10 mM iodoacetamide (IAA) for 30 min. The samples were digested with trypsin (Pierce; 75 ng/μl) for 18 h at RT. Peptides were labeled with tandem mass tag 18-plex (TMTpro) multiplex reagent (Thermo Fisher Scientific) following manufacturer’s protocol. SDC was precipitated with formic acid (FA) at final concentration of 2% (v/v) and centrifugation for 5 min at 10,000 rpm. Supernatant containing TMT-labeled peptides was dried.

### Mass spectrometry analysis

TMT-labeled peptides were fractionated using Waters XBridge C18 column (2.1 x 150 mm, 3.5 μm) and Dionex ultimate 3000 HPLC system. Mobile phase A consisted of 0.1% ammonium hydroxide; mobile phase B consisted of 100% acetonitrile and 0.1% ammonium hydroxide. Stepwise separation of peptides was achieved with a gradient elution at 200 μl/min: isocratic for 5 min at 5% phase B, gradient for 40 min to 35% phase B, gradient to 80% phase B in 5 min, isocratic for 5 min, and re-equilibrated to 5% phase B. Fractions were gathered in a 96-well plate every 42 sec. dried, and reconstituted in 50 μl 0.1% formic acid. The samples were analyzed on an Orbitrap Ascend mass spectrometer. A Dionex Ultimate 3000 system and mass spectrometer (Thermo Fisher Scientific) were used for data acquisition.

Analysis was performed as previously^26^ with some modifications. The samples were analyzed using a Real-Time Search-SPS-MS3 method. Ca. 3 µg of peptides / fraction were injected into a C18 column (Acclaim PepMap 100, 100 µm × 2 cm, 5 µm, 100 Å) at a 10 µl/min flow rate.

Separation was achieved via a 120-min low-pH gradient on a nanocapillary reversed-phase column (Acclaim PepMap C18, 75 µm × 50 cm, 2 µm, 100 Å) at 50°C. MS1 scans were collected over a 400-1600 m/z range using an Orbitrap at 120,000 resolution, with standard AGC and auto injection time, and included 2-6 precursor charge states. A dynamic exclusion window of 45 sec was applied with a repeat count of 1, mass tolerance of 10 ppm, and isotope exclusion permitted.

MS2 spectra were acquired in the ion trap at a turbo scan rate with HCD collision energy set to 32% and a maximum injection time of 35 ms. These spectra were explored versus the *Mus musculus* and CR proteomes using the Comet search engine. Static modifications included cysteine carbamidomethylation (+57.0215) and N-terminal/lysine TMTpro (+304.2071), with variable modifications including Asn/Gln deamidation (+0.984) and Met oxidation (+15.9949), allowing up to two variable modifications and a maximum of four peptides per protein. Precursors meeting these parameters were chose for SPS10-MS3 scans, performed at an Orbitrap resolution of 45,000 with normalized HCD collision energy set to 55%, AGC at 200%, and a maximum injection time of 200 ms.

### Proteomics data normalization, analysis, and visualization

To compare the results between sample batches, the datasets were normalized in various steps. For mouse proteins, normalization factor for each condition was calculated by the sum of all raw abundances in each TMT channel and dividing it by the maximum sum within given TMT-plex. For proteins with at least 60% of the TMT channels present, the remaining missing values were imputed using the minimum intensity observed within the corresponding TMT batch. To correct for potential batch effects, the data were scaled within each biological group. PCA analysis was performed using log_2_(scaled abundance) of proteins (Table S5) in the Perseus software version 2.0.11^64^. Log_2_ fold changes (log_2_FC) were calculated by comparing CR-infected samples to their corresponding UI controls. Differential protein expression between CR_WT_ and CR_M12_ was assessed using two-sample t-tests (p < 0.05) in Perseus. Two-sample t-tests were also performed for differential protein expression between CR_WT_ and UI, or CR_M12_ and UI. Proteins were considered significantly up- or down-regulated if they met the statistical threshold (p < 0.05) and a fold-change cutoff (|log2FC| ≥ 0.5). For data visualization, heatmaps and hierarchical clustering, the online Phantasus tool version v1.25.4 was used^65^. Volcano graph was prepared using VolcaNoseR online tool^66^.

### Fecal sample ELISA

Fecal samples were weighed, suspended in 1 ml PBS/0.1% Triton X-100 per 100 mg and homogenized for ∼ 15 min. Following brief centrifugation the supernatant was stored at -80°C. The concentrations of LCN2/NGAL and S100A8 was determined using DuoSet mouse Lipocalin-2 and S100A8 ELISA (R&D systems; Table S4) according to the manufacturer’s recommendations. Colorimetric readings were obtained using a FLUOstar Omega microplate reader (BMG biotech).

### Explant cytokines and chemokines

0.5 cm distal colon were weighed, immersed in RPMI medium with 100 μg/ml streptomycin and 100 U/ml penicillin, and incubated for 2 h at RT. The tissue was washed in complete RPMI media (containing 10 mM HEPES, 1 mM sodium pyruvate, 10% FBS and 100 μg/ml streptomycin and 100 U/ml penicillin and was assessed), followed by culturing in the same medium for 24 h at 37 °C, 5% CO_2_. The supernatant was centrifuged for 20 min at 3000 x g at RT and stored at -80 °C. The level of cytokine and chemokine was determined according the the manufacturer’s protocol. The panel included IL-22, Il-17A, IFN-γ, IL-1β, TNF, IL-18, CXCL1, IL-6, IL-10, CCL3, IL-23, G-CSF, and GM-CSF. Cytokine and chemokine levels were acquired using a FACSCalibur flow cytometer (BD Biosciences). The analysis was done using LEGENDplex data analysis software (BioLegend). Cytokines which showed values below the limit of detection in 90% of the samples were excluded.

### Isolation of lamina propria immune cells from mouse colon

Antibodies and blocking agents used for flow cytometry are listed in Table S4. Lamina propria cells were isolated from mice colons as previously described^67^. The 3.5 cm distal colons from infected or UI mice were excised, cleaned, cut opened longitudinally, and then incubated for 20 min at 37 °C in a shaking incubator in 1X HBSS (Ca^2+^ and Mg^2+^ free) supplemented with 2% FBS, 10 mM EDTA, and 1mM DTT. The cell suspension was centrifuged to separate lamina propria from IECs. Supernatants containing IECs were separated, and residual tissues were digested by incubating them at 37 °C for 40-50 min in RPMI 1640 media supplemented with 62.5 μg/ml Liberase, 50 μg/ml DNAse I (Sigma-Aldrich), and 2% FBS, passed through 100 μM cell strainer to obtain single cells.

### Flow cytometry

To stain for extracellular markers, single-cell suspensions or stimulated cells were plated in a 96 well V bottomed plate and stained for 10 min with LIVE/DEAD fixable blue (diluted in D- PBS) to detect and exclude dead cells from subsequent analysis. Cells were then treated for 20 min with Fcγ receptor block (BD Biosciences) followed by surface marker staining using fluorophore-conjugated monoclonal antibodies. All incubations were performed at 4°C in the dark unless otherwise stated. Negative (unstained and live/dead) along with fluorescent-minus-one (FMO) controls were considered to estimate background fluorescence. Cells were then washed and fixed for 20 min with using the eBioscience Forkhead box protein 3 (Foxp3)/transcription factor fixation buffer set. Fixed cells were kept in the dark at 4 °C until analysis.

Single stain controls for compensation were prepared using UltraComp eBeadsTM Compensation Beads. Cells were then washed prior to flow cytometry analysis on 50,000 live cells on an Aurora flow cytometer (Cytek Biosciences). Flow cytometry data were analyzed using FlowJo software (v10.8.1, Tree Star).

In the CD45^+^ population of live and single cells, neutrophils were defined as CD11b^+^ Ly6G^+^ cells, monocytes as CD11b^+^ CD11c^-^ F4/80^-^ Ly6C^-^ or iMonocytes (Ly6C^+^) cells, macrophages as CD11b^+^ CD11c^+^ MHCII^+^ F4/80^+^ CD64^+^, cDCs as CD64^-^ MHCII^+^ CD11c^+^ Ly6c^-^, pDCs as CD11b^-^ CD11c^+^ F4/80^-^ Ly6C^+^ cells, B cells as B220^+^ CD3^-^, and T cells as B220^-^ CD3^+^ (either CD4^+^ or CD8^+^) (gating strategy shown in Fig. S8).

### Statistics

Investigates were designed as randomized block of 4-5 mice per group per experiment; each experiment was repeated at least twice, unless specified. The number of mice used for each experiment is listed in Table S2. Results from all mouse investigations were pooled and analyzed. Flow cytometry data were analyzed using FlowJo software (v10.8.1, Tree Star). Two technical replicates were used for ELISA to determine the experimental mean. Statistical significance between two normally distributed groups was assessed by two-tailed t test, and that of more than two groups was assessed by analysis of variance (ANOVA) with indicated post-test. False discover rate (FDR; Q=5%) was used to correct for multiple comparisons (Tukey’s or Sidak’s correction), or as implemented by GraphPad Prism 10.4.0. Data plotting and statistical analysis were performed using Prism 10.4.0 (GraphPad Software Inc.). Statistical details of experiments are described in the figure legends. Custom images in Fig. 1, 2, and S1 were made using BioRender (using biorender.com).

## Acknowledgements

We thank Prof. Brigitta Stockinger, The Francis Crick Institute, for providing the C57BL/6 *Il22^-/-^* mice used to establish the colony at Imperial College London. We thank Jessica Rowley and Larissa Zarate Garcia for their technical assistance with Flow Cytometry. The work was supported by Wellcome Trust Investigator Award grants 107057/Z/15/Z and 224282/Z/21/Z, and MRC programme grant MR/R02671, and grant RYC2021-031342-I funded by MICIU/AEI/10.13039/501100011033 and by European Union Next Generation EU/PRTR.

## Author notes

All authors declare no conflict of interest.

## Data availability statement

Identifier for the PRIDE dataset: PXD062886

**Password:** x8cjWgY9y8rr

The supporting the finding of this study are available as follows:

Table S2: Mice numbers

Table S3: Bacterial strains, plasmids, and primers

Table S4: Key resources

Table S5. Proteomics of mouse proteins of uninfected mice and following infection with CR_WT_ or CR_M12_

Key resources data: raw measurements shown in all the main and supplementary figures

## Generative Artificial Intelligence

We declare the usage of Chat GPT-4o model for improving Grammer, readability and clarity of text.

